# RNA structures and dynamics with Å resolution revealed by x-ray free electron lasers

**DOI:** 10.1101/2023.05.24.541763

**Authors:** Kara A. Zielinski, Shuo Sui, Suzette A. Pabit, Daniel A. Rivera, Tong Wang, Qingyue Hu, Maithri M. Kashipathy, Stella Lisova, Chris B. Schaffer, Valerio Mariani, Mark S. Hunter, Christopher Kupitz, Frank R. Moss, Frédéric P. Poitevin, Thomas D. Grant, Lois Pollack

**Affiliations:** School of Applied and Engineering Physics, Cornell University; Ithaca NY 14853 USA; Meinig School of Biomedical Engineering, Cornell University; Ithaca NY 14853 USA; Linac Coherent Light Source, SLAC National Accelerator Laboratory; Menlo Park, CA 94025 USA; Department of Structural Biology, Jacobs School of Medicine and Biological Sciences; University at Buffalo, Buffalo, NY 14203 USA

## Abstract

RNA macromolecules, like proteins, fold to assume shapes that are intimately connected to their broadly recognized biological functions; however, because of their high charge and dynamic nature, RNA structures are far more challenging to determine. We introduce an approach that exploits the high brilliance of x-ray free electron laser sources to reveal the formation and ready identification of Å scale features in structured and unstructured RNAs. New structural signatures of RNA secondary and tertiary structures are identified through wide angle solution scattering experiments. With millisecond time resolution, we observe an RNA fold from a dynamically varying single strand through a base paired intermediate to assume a triple helix conformation. While the backbone orchestrates the folding, the final structure is locked in by base stacking. In addition to understanding how RNA triplexes form and thereby function as dynamic signaling elements, this new method can vastly increase the rate of structure determination for these biologically essential, but mostly uncharacterized macromolecules.

## Introduction

The majority of genomic DNA is transcribed into RNA, yet only a small fraction is translated into proteins (*1*). Although the biological roles of much of this untranslated RNA have yet to be elucidated, non-coding RNA is increasingly linked to vital cellular functions (*2*) and has great potential as a therapeutic (*3*). Significant advances in assigning biological roles to these transcripts, or in improving the targeted design of drugs, may be accelerated if their structures can be deduced. Macromolecular structures can be directly measured, simulated, or derived from sequence by solving the folding problem (*4*), using computational tools (*5*), or applying artificial intelligence (*6*). Because of the highly charged and dynamic nature of its backbone, as well as the similarity of its nucleotide building blocks, RNA structures are far more challenging to solve than protein structures, by either experiment (*7-9*) or all atom simulation (*10, 11*) The paucity of measured structures limits our ability to advance the field (*12*), especially when compared to the state-of-the-art in protein structure prediction (*6*).

X-ray scattering from biomolecules in solution has great potential to reveal their full structure, from overall size down to Å dimensions. Small angle X-ray scattering (SAXS) has been frequently applied to measure RNA structure (*13-15*) and informs about global, nanometer level molecular structures (*16*). Scattering to large angles (wide angle X-ray scattering, WAXS), enhances measurement resolution to the single Å (*17, 18*). Recent WAXS studies (*19-23*) have begun to connect distinctive peaks in scattering profiles with real space features of DNA and RNA, including backbone geometries, structural building blocks (duplexes), and base stacking. However, because WAXS signals from biomolecules are typically 100-1000 times smaller than SAXS signals, access to this information rich, higher resolution region requires seconds-long exposures of typical millimeter or sub-millimeter sized solution samples with volumes of 10 ‘s of microliters, even at high flux synchrotron sources (*19*). Although these measurements hold promise for expanding the database of RNA structures, visualization of folding reactions would enhance our ability to predict structure from sequence offering a general approach. Rapid fluidic mixers are required to measure RNA folding on the relevant, millisecond time scales (*24*). X-ray illuminated sample volumes within these flow cells can be 100 to 1000 times smaller than used for equilibrium measurement (*25*), depending on the desired time resolution. These time-resolved WAXS measurements are beyond the current-state-of-the-art at synchrotrons.

X-ray free electron laser (XFEL) sources present a unique opportunity to illuminate micron scale, 10 ‘s of femtoliter sample volumes with high flux beams. At the Linac Coherent Light Source (LCLS, SLAC), beam intensities at the sample are ∼1000 times higher than at state-of-the-art synchrotron beamlines, enabling measurement of previously impossible-to-detect signals from time-resolved WAXS (*26-29*). We dramatically expand this field by introducing mixing injectors (*30, 31*) to perform the first, to our knowledge, chemically triggered time-resolved solution scattering experiments on biomolecules at an XFEL. This experiment follows the dynamic acquisition of secondary and tertiary structure of an unstructured, single strand of RNA, as it folds to a triple helical final state. RNA triplexes are of particular recent interest and are gaining attention as functional motifs and drug targets (*32, 33*). Comparison with WAXS profiles of an RNA motif library, created by extending the angular range of scattering reported in prior studies, allows us to identify distinctive experimental signatures of single strand, duplex, and triplex molecular forms. These findings demonstrate the power of WAXS to reveal RNA structural motifs, to 3 Å resolution, in both static and dynamically folding samples. When coupled with SAXS, WAXS offers a new method for room temperature structure determination of nucleic acids, closing the gap between structure and prediction for these hard-to-characterize molecules (*12*).

## Results

### High resolution features in solution scattering profiles provide incisive information about RNA ‘s molecular state

The wide angle scattering regime, q > 0.5 Å^-1^ (q = 4πsinθ/λ, where θ is half the scattering angle and γ is the x-ray wavelength) is information-rich for nucleic acid samples; to the best of our knowledge RNA scattering profiles that extend beyond q = 1 Å^-1^ have never before been published. Figure 1, panels A-C shows measured scattering profiles of three different RNA motifs, along with a cartoon representation of the structures they reflect. These curves were acquired at synchrotrons. Panel A shows WAXS profiles of RNA single strands, both unstructured chains of 30 uracil nucleotides, rU30, and minimally structured chains of 30 adenosine nucleotides, rA30. Low angle scattering (q < 0.25 Å^-1^) profiles were previously published for single stranded RNA, and the structures shown below were derived from those measurements (*34*). Panel B shows the scattering profile of a designed duplex that terminates in a loop. Panel C shows scattering profiles of RNA triplexes with different lengths, constructed by adding a third, triplex forming strand to a hairpin duplex. Lower q portions of these data (for q < 1.0 Å^-1^) were previously published (*19*) and best-fit structures were determined by comparison with all atom simulations. Distinct features of each profile can be associated with structural patterns that characterize single, double, or triple-stranded RNA, highlighted by colored bars in panels B, C, E, and F. These connections are explained in more detail below and in Figure S1.

**Figure 1.**
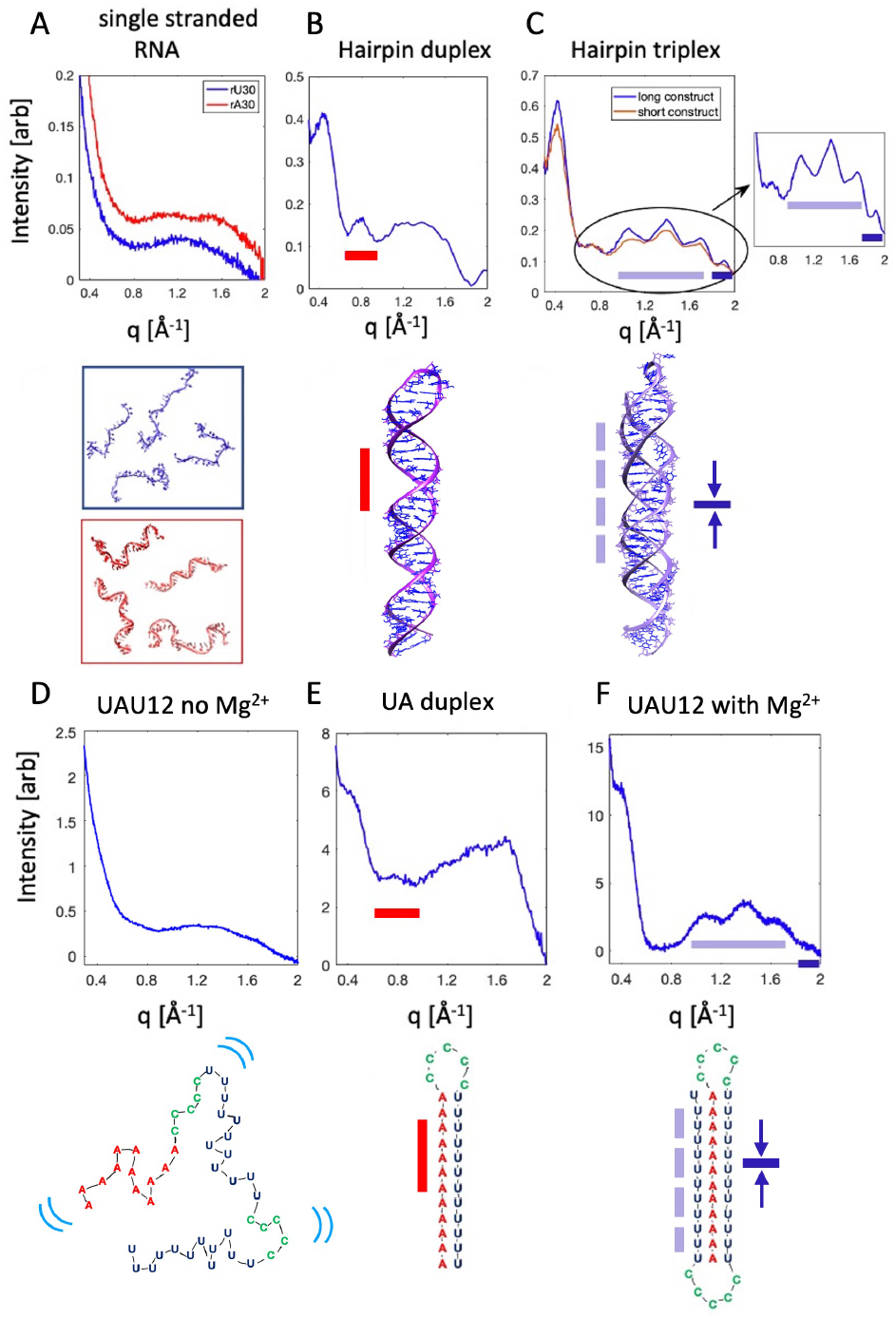
WAXS features associated with structural motifs of RNA. A) scattering profiles of single stranded constructs rU30 and rA30 shown as intensity vs. q. Representative structures in the experimentally determined ensemble of rU30 (top) and rA30 (bottom) are shown below. B) scattering profile of designed RNA hairpin duplex from Ref. (*19*), with a model of one duplex conformation shown below. C) scattering profile of RNA hairpin triplexes from Ref. (*19*) with a model of triplex from Ref. (*19*) shown below. The inset emphasizes the rich information content of the WAXS region. The colored lines in panels B and C link the features in the scattering profile with their corresponding real space structures. D) scattering profile of the t=0, Mg^2+^ free starting state of the time resolved experiment, acquired at LCLS, with a cartoon representation of unfolded UAU12. Blue lines indicate dynamic motion. E) scattering profile of designed hairpin duplex with UA base pairs, with a cartoon duplex shown below. F) scattering profile of folded UAU12 triplex, 1 second after the addition of Mg^2+^ to trigger folding, acquired at LCLS, with a cartoon model depicting the folded triplex shown below.

#### Single strands

Figure 1 panel A shows the measured scattering profile of two single stranded constructs rU30 and rA30 in buffered solutions containing 100 mM NaCl. Single stranded RNA is dynamic and highly flexible, thus their molecular structures are best recapitulated by an ensemble whose properties (and summed scattering profiles) are consistent with experimental measurements from a variety of probes (*34*). Typical structures from a best-fit ensemble are shown below the scattering profiles of panel A (*34*). The intensity of scattering from these disordered molecules decreases monotonically from low q to about q=0.8 Å^-1^, where a broad peak is visible for both rU30 and rA30. This peak occupies a range of q values corresponding to length scales between about 3 and 8 Å. As noted in prior work on DNA (*23*), well defined peaks in this range are derived from the phosphate backbone. In contrast, the broad peak seen here reflects the lack of structured elements on these length scales, hence the structural heterogeneity of the ensemble. Within it, signs of structure emerge for rA30, in the form of two loosely separated, broad peaks near q= 1.1 and 1.5 Å^-1^, but not for rU30, consistent with base stacking induced ordering present in the former and absent in the latter (*34*). Both curves decay at larger q.

#### Duplexes

Figure 1 panel B shows the scattering profile of a 62 nucleotide RNA hairpin, consisting of a 29 base-paired duplex, terminated by a four nucleotide loop from Ref. (*19*). Prior studies examined the scattering out to q =1.25 Å^-1^ and used molecular dynamics simulations (*19*) in conjunction with machine learning (*21*) to confirm that the small, but notable peak at q= 0.8 Å^-1^ is connected with the helical major groove, a repeated spatial dimension in the duplex structure. A red bar in the figure connects these reciprocal and real space features. At higher angle, 1 < q < 2 Å^-1^, a broad peak appears. Although better articulated than in panel A, the sharp peaks characteristic of DNA duplexes (*23*) are absent, suggesting that this duplex has significant conformational variation. This observation is consistent with other studies of RNA duplexes (*20, 22*). A single molecular conformation from the simulations of Ref. (*19*) is pictured just below the profile.

#### Triplexes

Figure 1 panel C illustrates the unique and never-before-reported triplet of peaks associated with triple helical backbones, indicated by a purple bar in the figure. Measured scattering profiles from two triplexes are shown. Both were constructed by adding a triplex forming single strand to a hairpin duplex. The shorter/longer construct consists of loop terminated 17/29 base paired duplexes, with an added 12/24 nucleotide single strand that binds to form base triples over a significant length of the molecule. A model of this latter structure is shown just below the plot, taken from Ref. (*19*). Its increased length reinforces the features that uniquely identify the triplex structure by WAXS: three sharp peaks at q = 1.0, 1.4 and 1.7 Å^-1^. By analogy with the DNA fingerprinting studies of Ref. (*23*), and structural modeling (SI: *Modeling connects features in the scattering profile with molecular structures* and Figure S1), these peaks reflect the regular spacing of atoms along the backbone in a helical conformation. This cluster of three peaks is flanked by additional peaks on either side. At lower q, a weak reflection of the major groove peak can be seen. Figure S2 shows that this peak is significantly reduced and shifted to lower q between the duplex and triplex states. As previous x-ray fiber diffraction studies of a pure RNA triplex suggest that it is roughly cylindrical, lacking grooves (*35*), this reduced peak may reflect a small amount of duplex that is present in the construct shown below panel C. At higher q (1.93 Å^-1^), the small but distinct peak (dark blue bar, Figure 1C) reflects the 3.26 Å distance between stacked base triples (Figure S1and Ref. (*19*)). The sharpness of the peaks indicates that triplex molecular dimensions are well defined, with less conformational variation than either the duplex or single strands.

These curves reveal five distinct features of WAXS (q > 0.5 Å^-1^) scattering profiles of RNA structures, only one of which has been previously identified (*19*). The peak at q= 0.8 Å^-1^ reflects the existence of a major groove. The three peaks at q= 1.0, 1.4 and 1.7 Å^-1^ represent regularly positioned atoms in the three RNA backbones of the triplex and the tiny peak at q= 1.9 Å^-1^ reflects stacking of base triples (Figure S1 and length scales extrapolated from structures of Ref. (*19*)). Additional information can be gleaned from lower q, SAXS features (SI: *Low q changes* and Figure S2). In particular, when unstructured single strands combine to form a duplex, a peak appears near q = 0.4 Å^-1^. According to machine learning models (*21*), the position of this peak reflects the radius of the helical structure. It shifts slightly to lower q for the triplex relative to the duplex, reflecting the larger helical radius of the former. Although this work highlights higher angle features, the lower q peak also contributes to our understanding of RNA structure and serves as an important milestone in folding studies.

#### Starting and ending states of the mixing experiment

For folding studies, we used a 46 nucleotide RNA with sequence 12U-5C-12U-5C-12A. This construct is a long single strand of RNA, in contrast to the triplexes shown in panel C, which were each created from two separate RNA molecules. This design is based on a well-characterized triplex forming sequence (*36, 37*). We refer to it as UAU12, to highlight the capture of the A12 strand between the two U12 strands. Measured scattering profiles of the starting and ending states of the folding experiment are shown in Figure 1, panels D and F. These data were acquired at the XFEL. In the initial, unfolded state the RNA is in a low ionic strength buffer. Folding is initiated by the addition of MgCl^2^.

Structural details about the RNA during the time resolved experiment can be gleaned by comparison with the profiles of the reference systems in panels A-C. For example, the similarity of the scattering profiles of panels D and A suggests that the initial state of the time resolved experiment is unstructured. This assessment is based on the absence of distinct features in its scattering profile. A cartoon that suggests this conformation is shown below the profile of panel D; here, the blue lines indicate dynamics.

Figure 1 panel E shows the measured scattering profile of a related hairpin duplex (12U-5C-12A, the first 29 nucleotides of the full UAU12 construct), folded by adding Mg^2+^ to the solution of single strands of this construct. This profile was acquired to identify a potential folding intermediate, a 12 base pair A-U duplex. Because RNA duplexes are length (*20*), sequence (*38*) and salt (*11*) dependent, it was critical to measure the structure of a molecule where an rA12 strand binds to an rU12 strand to form a 12 base pair duplex. The duplex of panel B is much longer, and of mixed sequence, hence will have better defined structures. Collections of rA and rU strands tend to form triplexes when combined, in addition to duplexes (*39*), but the mixed sequence 29 nucleotide strands formed duplexes: the scattering profile of panel E displays comparable features to the duplex profile of panel B. A peak detected near q=0.8 Å^-1^, indicated by the red bar, reflects the formation of a duplex major groove (*21, 22*).

Finally, the ending state of the experiment (UAU12 with added Mg^2+^), shown in panel F, displays the characteristic three-peak structure of a triplex (purple bar), as well as the higher q peak of panel C (blue bar). The major groove peak (at q=0.8^-1^) is absent from this measured profile, consistent with third strand filling the major groove. The cartoon schematically indicates the conformation of the folded triplex.

### Mixing injectors target time scales from the single millisecond to the single second

To capture WAXS profiles with sensitivity to millisecond scale conformational changes, time-resolved data were collected from micron-sized liquid jets at the CXI beamline at LCLS. Mixing injectors (schematically shown in Figure 2) were used to initiate the Mg^2+^ mediated folding of UAU12. A full description of the fabrication and operation of the injectors, including design parameters, can be found in Ref. (*30*). Three different injector geometries were used to access the broad range of time points of interest to this folding experiment: t= 0, 6, 10, 60, 100, 500 and 1000 milliseconds. By varying the flow rates of the sample and buffer, each injector can be used to acquire data at multiple, closely spaced time points. On average, about 16,000 good quality single shot profiles are used to generate the averaged profile required to visualize the WAXS features from RNA flowing in a sheathed, micron-sized jet. With the current XFEL repetition rate of 120 Hz, it takes just over 2 minutes of beam exposure to acquire such a profile. Given that a data point requires both a sample-present (RNA in buffer) and a sample absent (buffer alone) exposure, a total of 4 minutes of data acquisition yields one time point. Details about profile acquisition, computation, normalization, selection, and background subtraction are provided in Methods and Figures S4-S8.

**Figure 2:**
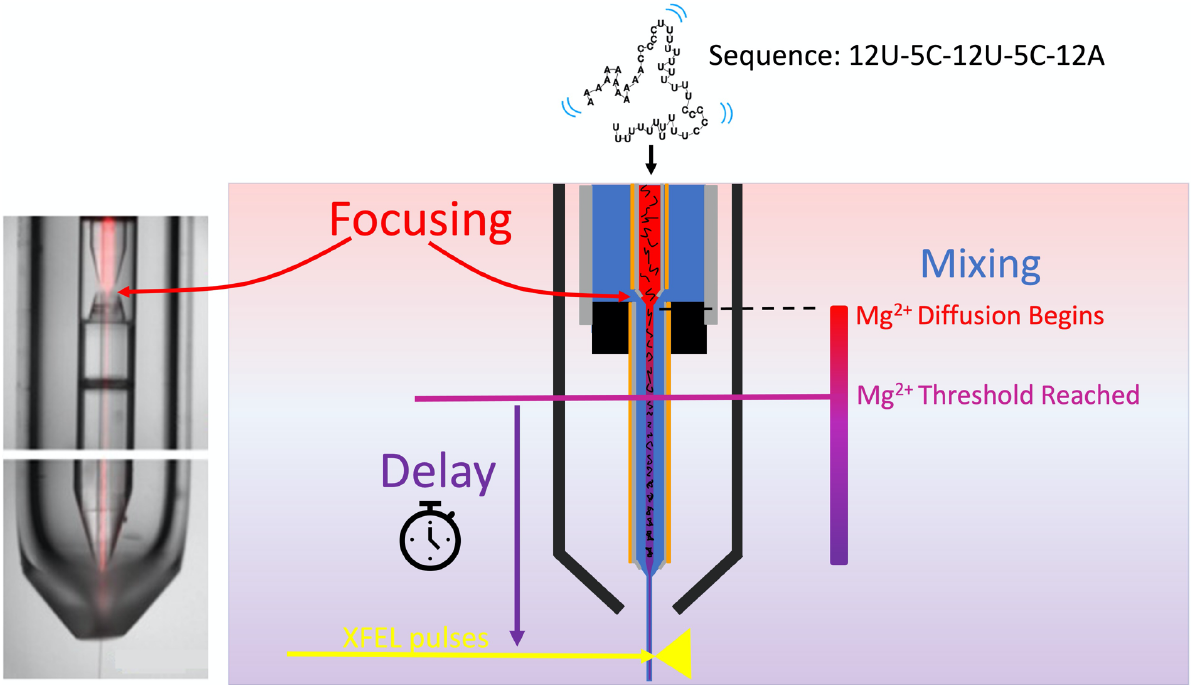
Mixing injectors for time resolved studies at the XFEL. A solution containing RNA in low ionic strength buffer (red liquid) is hydrodynamically focused into a thin cylindrical stream, which is surrounded/sheathed by a buffer containing additional Mg^2+^ ions (blue liquid). These streams flow coaxially through a constricted region, where mixing occurs by the rapid diffusion of Mg^2+^ ions into the thin, central, RNA-containing stream. The reaction is initiated once the [Mg] reaches a carefully selected threshold (Figure S3). The reported time point for each measurement corresponds to the time it takes for the molecule to flow to reach the beam. This delay time is carefully determined for each experiment. Detailed parameters for injectors can be found in Methods and Tables S1-2.

### Time resolved studies reveal folding intermediates; a transient duplex forms before the triplex

Time-resolved experiments were performed at RNA concentrations of 1 mM and 0.5 mM. Different amounts of Mg^2+^ were required to quickly reach the reaction initiation threshold for each [RNA] (SI: *Determining the proper [Mg*^*2+*^*] level to initiate the* reaction and Figure S3). Figure 3 shows the time series for the higher concentration experiment. These curves show the time progression of scattering profiles from the form shown in panel 1D to that of panel 1F. For this series, we acquired good quality data at t= 6 ms, 10 ms, 60 ms and 1000 ms. The curves are shown along with the t=0 state (acquired at [RNA]=0.5 mM). Four features are visible at the earliest time point acquired, 6 ms after the initiation of folding (orange). A shoulder appears near q=0.4 Å^-1^ and the major-groove associated peak appears near q = 0.8 Å^-1^. Two higher q peaks also appear, though they are not well resolved. On this short time scale, the lower q data indicate that some fraction of the sample is in the duplex state. The appearance of triplex-associated peaks also suggests partial, but not yet full structuring of the triple helical backbone. It is unclear whether all molecules fold through a mandatory duplex intermediate, or whether some fold directly to triplex. This distinction will require measurements at shorter time points. Nonetheless, it is clear that the RNA shows some duplex and some weak triplex features within 6 ms of folding initiation.

**Figure 3.**
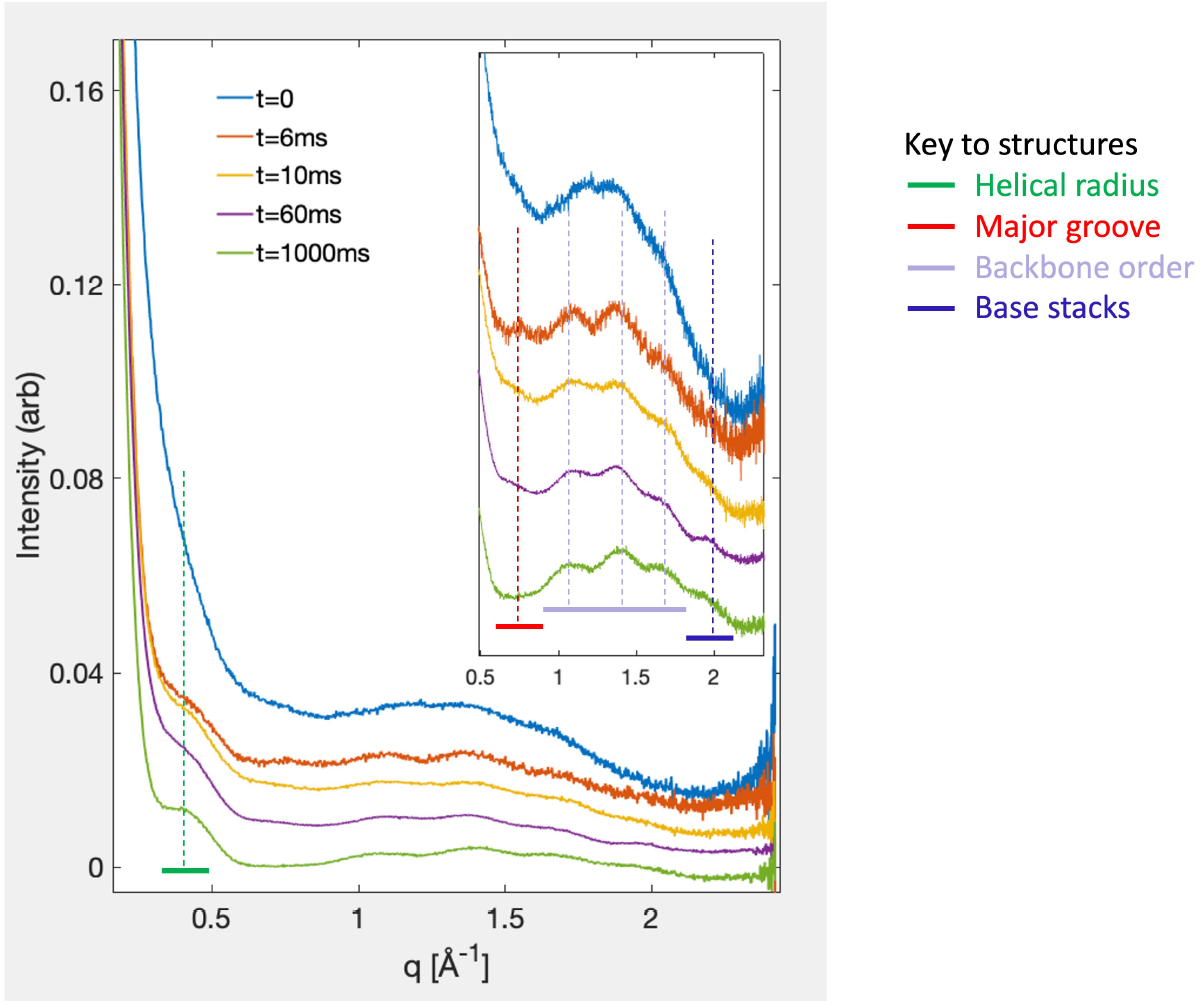
Time resolved WAXS data following the formation of an RNA triplex from a minimally structured single strand. The measured time series following folding of the UAU12 construct from single strand to triplex is shown for the initial (t=0) and final folded (t=1000 ms) states, along with three intermediate time points. Data were acquired 6, 10 and 60 ms after folding initiation. Distinct features in the scattering profiles, identified in conjunction with equilibrium measurements, modeling and machine learning studies shown in Figure 1, aid in the interpretation of the dynamically evolving structures that populate the folding landscape. Features of disordered, partially ordered and fully ordered RNA are observed as the reaction proceeds. Together, they reveal the folding strategy of this molecule.

In the 10 ms profile (yellow), diminished duplex features are observed: the major groove peak is reduced. The triplex peaks are better articulated, and a small, but not quite resolvable, perturbation in intensity appears near the highest q, base stacking peak. Sixty ms after the initiation of folding (purple curve), the triplex peaks, as well as the base stacking peak are firmly established. These trends are further underscored at 1000 ms after folding (green). Here, the lower q shoulder near q=0.4 Å^-1^ has settled into a form that resembles that of the static triplex (Figure S2). At this time point, the scattering profile displays all of the SAXS/WAXS peaks seen in an equilibrium, folded curve, acquired on our lab source (Figure S9). Deviations in the baseline levels can be explained by temperature variations (SI: *The effect of temperature on scattering profiles of nucleic acids* and Figures S10-S11).

## Discussion

Overall, the RNA ‘s folding strategy can be established directly from the curves shown in Figure 3. The triplex folds from a mostly unstructured single strand, through a duplex intermediate, to a state where the three backbone strands form the outline of the triplex. Finally, base stacking appears at longer times, locking the molecule into its final structure. This strategy is suggested in Figure 4.

**Figure 4.**
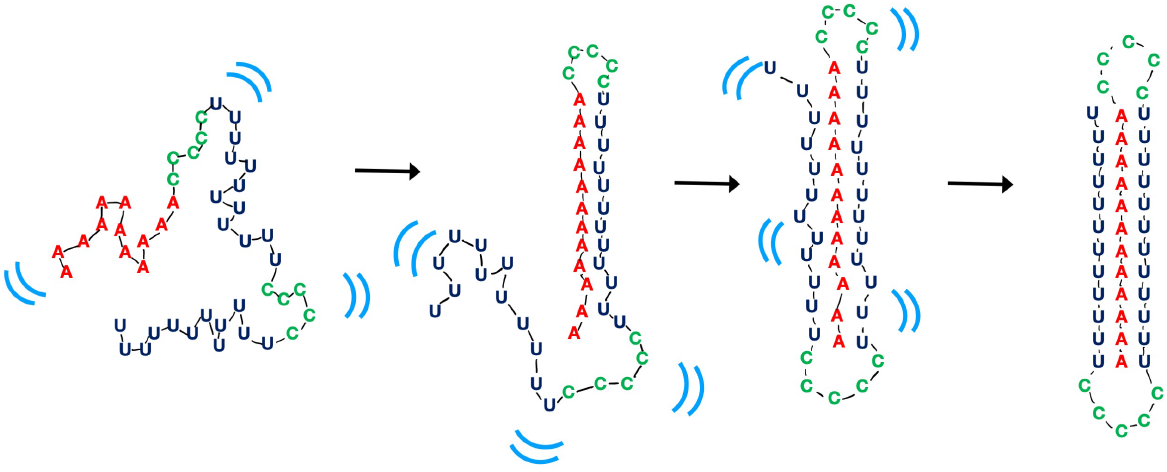
Schematic cartoon depicting folding dynamics from single strand to triplex as read directly off the scattering profiles. The left-most cartoon depicts the dynamic variation of the low ionic strength starting state: a mostly unstructured and dynamic (indicated by blue marks) single strand of 46 nucleotides. The next cartoon is a representation of a transiently detected state in which one 12 base paired A-U duplex forms. The duplex is identified by the ‘major groove feature ‘ that appears in the t=6 ms scattering profile. The third panel represents a state in which the backbones have formed a loose triplex structure, defined by the appearance of a triplet of peaks at higher q. The final structure, at right, appears to have a less dynamic structure, and shows strong evidence of stacking of base triples that lock the molecule into a more rigid structure. The sharp peaks of the WAXS profile suggest that the molecule is less dynamic in this state; the triplex structure is well-defined with much less variation than RNA duplex structure.

The order of appearance of all of the peaks is recapitulated by the measured time series at [RNA]=0.5 mM, where data points were acquired at 0, 6, 100 and 500 ms (SI: *RNA folding at reduced concentration: 0*.*5 mM* and Figure S12). Most significantly, the transient duplex intermediate again appears at 6 ms, and vanishes at all later times. Slight deviations were seen in the amount of triplex present at 6 ms, likely a result of mixing conditions.

This distinctive view of dynamic structural changes from single strand to triplex underscores the highly dynamic nature of the RNA backbone, even in the duplex state, until it captures a third strand in the proper triplex geometry. Once the proper backbone arrangement is assumed, the structure is locked into place by base stacking. This experiment provides direct evidence that the RNA backbone drives folding, a conclusion that was previously hypothesized by comparison with MD simulations (*24*), but not directly observed until now.

In summary, WAXS reveals numerous features of secondary and tertiary RNA structures, which are instantly identifiable as the data are acquired. Our studies reveal the dynamic appearance and disappearance of helical grooves, backbone ordering and base stacking. This structurally detailed information is trivially extracted by inspection of profiles of equilibrium (static) states, as well as from transient states during a folding reaction. When coupled with lower angle data that reveal the relative spatial arrangement of these structural elements, this work highlights the potential of solution scattering to uniquely and directly characterize both static and dynamic RNA structures.

## Conclusions

Presently, less than 2000 RNA structures are deposited in the protein data bank. If a substantial database of structures can be built, either directly or by solving the folding problem, new deep learning techniques (*12*) can potentially provide a structural revolution in the RNA world comparable the one recently provided for proteins by Alpha Fold (*6*). High angle solution x-ray scattering provides an RNA structural motif library to easily link distinct features in scattering profiles with real space RNA structures. When coupled with lower angle data that articulate the relative arrangement(s) of these motifs, this technique can help address this unmet need to solve RNA structures.

In addition to enhancing structure solving methods, this work highlights the unique ability of highly brilliant, XFEL x-rays to provide the exceptional sensitivity needed to observe subtle structural changes in macromolecular systems. Although there is already much to learn from this now demonstrated technique, XFEL technology is rapidly advancing. The increasing pulse repetition rates at XFELs worldwide (from hundreds to millions of pulses per second (*40*)), coupled with the development of fast framing detectors to handle these data rates, will enable the visualization of even higher q (sharper spatial resolution) features in solution scattering. Once these upgrades are realized, atomic resolution in solution structures, at room temperature, and on biologically relevant time scales is imminent.

## Materials and Methods

### Nucleic acid samples

All RNA and DNA samples were purchased from Integrated DNA Technologies (Coralville, IA) as single strands. Molecular reconstitution, including buffers and annealing protocols are distinct for each sample, and are described in detail in SI: *Materials and Methods*.

### Mixing Injectors

Mixing injectors (as described in (*30, 41*)) were designed and fabricated for the time-resolved experiments. These devices utilize a flow-focused diffusive mixer coupled to a Gas Dynamic Virtual Nozzle (GDVN) to initiate reactions just prior to producing a freely flowing liquid jet for data collection. Design parameters, fabrication details, and specifications for each mixer used are described in SI: *Materials and Methods*.

### Data collection and analysis at XFEL

Time resolved RNA folding data, as well as data on the AT duplex were collected at the Coherent X-ray Imaging (CXI) instrument at the LCLS (*42*) (SLAC National Accelerator Lab, Menlo Park, CA), using the 1 micron sample environment. The mixing nozzles were loaded into the vacuum chamber on a nozzle rod (Standard Configuration 1). X-rays were delivered at 120 Hz frequency with a pulse energy of 2 mJ and beam size of ∼1 μm (FWHM). Scattering was collected on the Jungfrau-4M detector (*43*). The X-ray energy was 6 keV and the detector was positioned 106 mm from the sample resulting in a q range of 0.12 – 2.4 Å^-1^. Calibration of the detector distance and geometry was performed using silver behenate. Newly developed protocols for acquiring fully background subtracted solution scattering data at XFELs are provided in SI: *Materials and Methods*.

### Data collection and analysis at NSLS II

Solution X-ray scattering measurements on single stranded rU30 and rA30 RNA were performed at the 16-ID Life Science X-ray Scattering beamline at the National Synchrotron Light Source II (NSLS-II) of Brookhaven National Laboratory (*44*). Scattering from the small-angle (q=0.01-0.3Å^-1^) and wide-angle (q=0.3-3.2Å^-1^) regimes were read simultaneously using two Pilatus 1M detectors (Dectris, Switzerland, EU) arranged in series. The transmitted X-ray beam intensity was also recorded during each measurement. Centering and calibration of the beam on both detectors was performed using a silver behenate standard in BioXTAS RAW, as well as masking, radial averaging and buffer subtraction (*45*). Previously published data (*19*), reproduced here in Figure 1 panels B and C, were also acquired at NSLS II on the LiX beamline.

### Data collection and analysis on lab source

Measurements on the rUA duplex hairpin, as well as on the temperature dependence of the DNA AT duplex were performed using a BioXolver with Genics source (Xenocs, Holyoke, MA) using the setting WAXS_STD. Data collected from the BioXolver are analyzed using BioXTAS RAW, as described above. The quoted sample temperatures, when noted, were achieved by temperature controlling the sample capillary.

### Statistical Analysis

Time-resolved data were acquired from an average of 15,878 traces for each RNA containing profile, and 15,337 traces for each buffer background. At a frame rate of 120 per second, this corresponds to just over two minutes of data acquisition for each condition.

## Acknowledgments

We acknowledge Weiwei He, Yen-Lin Chen and Serdal Kirmizialtin for providing pdb files for the previously published hairpin duplex and triplex structures shown in Figure 1. We also acknowledge useful discussions with Rick Kirian about solution scattering at XFELs. We thank Shirish Chodankar for assistance with data acquisition at NSLS II. †Present address: Pfizer Inc. Pearl River, NY, ‡present address: Analytical Research and Development, Merck & Co., Inc., West Point, PA §present address: Altos Labs, Redwood City, CA.

## Funding

National Science Foundation through the BioXFEL STC award number 1231306 (LP)

National Science Foundation grant DBI-1930046 (LP)

National Institutes of Health grant R35-GM122514 (LP)

National Institutes of Health grant R01-GM133998 (TDG)

Natural Sciences and Engineering Research Council of Canada (NSERC) (TW)

U.S. Department of Energy, Office of Science, Office of Basic Energy Sciences under Contract No. DE-AC02-76SF00515 (use of the Linac Coherent Light Source (LCLS), SLAC National Accelerator Laboratory)

National Institutes of Health grant P41-GM139687 (LCLS/MSH)

National Institutes of Health grant P30 GM133893, S10 OD012331, and BER-BO 070 (for work performed at the CBMS beam line LIX (16ID) at NSLS-II)

U.S. Department of Energy grant BES-FWP-PS001 (NSLS II)

National Institutes of Health grant S10 OD028617 (LP BioXolver X-ray Source) National Institutes of Health grant S10 OD025079 (LCLS Jungfrau Detector)

## Author Contributions

K.A.Z. designed, fabricated and operated the mixing injectors and assisted in data acquisition, sample preparation and writing the manuscript. S.S. characterized the UAU12 construct and the reaction conditions, and assisted in LCLS experiments measuring the dA-dT duplex. S.A.P. acquired data on the UAU12 constructs and the dA-dT duplexes on the BioXolver source and prepared sample for LCLS experiments. D.A.R. and C.B.S. fabricated critical components of the mixing injectors. T.W. acquired WAXS data on the single strand constructs using NSLS II. Q.H. acquired SWAXS data on the hairpin duplex using the BioXolver source. M.M.K. assisted with sample preparation during the beamtimes. S.L. assisted with injector operation during beamtimes. V.M. helped specialize data analysis software for this experiment. C.K. assisted with sample preparation and injector operation during beamtimes. M.S.H., F.M. and F.P.P. operated the CXI station during LCLS beamtimes. T.D.G. assisted in real-time data visualization and creating and modifying data analysis software during and after the beamtime. L.P. assisted in data acquisition, analyzed the data, wrote the manuscript, supervised and coordinated all aspects of the experiment.

## Data and materials availability

Raw data for all scattering profiles of RNA and DNA are provided in the supplementary materials or in the SASBDB as indicated.

## Supplementary Materials

Supplementary text, figures, tables are available on-line.

