## Supporting Information for "RNA structures and dynamics with Å resolution revealed by x-ray free electron lasers"

Kara A. Zielinski *et al.*

**This PDF file includes:**

Supporting text
Figures S1 to S12
Tables S1 to S2

**Supporting Information Text**

### Materials and Methods

#### Sample Preparation

*Single stranded RNA sample preparation*
RNA samples were prepared as described in (*34*). Briefly, 30 nucleotide poly-uridine (rU30) and poly-adenine (rA30) RNA were purchased from Integrated DNA Technologies (Coralville, IA, USA), reconstituted in a storage buffer of 20 mM NaCl, 1 mM MOPS pH 7.0, 10 µm EDTA, and subsequently desalted and buffer exchanged into 100 mM NaCl, 1 mM MOPS pH 7.0, 10 µm EDTA for data collection. For samples containing magnesium, a small volume of 1M $MgCl_2$ was spiked into the sample shortly before the measurement. RNA concentrations of 600, 200, 50 µM were prepared, with the highest concentration aimed toward obtaining sufficient wide-angle scattering signal. All RNA samples were annealed by heating to 90°C for 3 minutes and snap cooling at 4°C for 20 minutes before measurements. Molecular structures were reproduced from SASBDB files: SASDFB9 for rA30 and SASDFK9 for rU30.

*Hairpin duplex and hairpin triplex preparation*
These scattering profiles were reproduced from Reference (*19*) and the SASBDB entries SASDKX5 for the hairpin duplex in 5 mM $Mg^{2+}$, and SASDK26 for the hairpin triplex in 5 mM $Mg^{2+}$. Molecules were prepared as described in that publication; no new profiles were measured for this work.

*A-U duplex preparation*
The 29 nucleotide construct U12-C5-A12 was purchased from Integrated DNA Technologies (IDT) and reconstituted in low salt buffer (1mM NaMOPS, 50uM EDTA and 2mM NaCl). It was buffer exchanged into a $Mg^{2+}$ containing buffer (1mM NaMOPS, 50uM EDTA, 2mM NaCl and 2.5 mM MgCl2) three times using 3k MWCO spin column at 14k xg for 10min at 4°C. The sample was heated to 85°C for 3 mins and cooled on ice. Data shown in Figure 1 were acquired on a sample with [RNA]=1mM, at T=10˚C; lower concentration samples acquired at room temperature show identical features in their scattering profiles.

*Triplex construct preparation*
The RNA construct, UAU12, with sequence 12U-5C-12U-5C-12A was purchased as a lyophilized powder from Integrated DNA Technologies, Inc. (Coralville, IA). The powder was reconstituted in a low salt buffer (1 mM NaMOPS, 2 mM NaCl, 50 µM EDTA at pH 7) for 15 minutes to maximize the yield. The sample was then buffer exchanged six times using a 10 kDa cutoff filter by spinning at 3600 G for 15 minutes at 4˚C to remove any impurities. A thermal heat block was used to anneal the sample by heating it to 92-94˚C for 5 minutes. The sample was removed from the heat and cooled to room temperature in about 15 minutes. Finally, the sample concentration was measured by UV-Vis absorbance and adjusted to either 1 mM or 0.5 mM for data collection. During all sample handling, gloves were worn, and other precautions were taken to prevent RNase contamination.

*A-T duplex preparation*

AT duplexes were comprised by base pairing dT25 with dA25. Both strands were purchased from Integrated DNA Technologies, Inc. (Coralville, IA). They were annealed at 95˚C for 5 minutes and slowly cooled to 22 ˚C in an hour. This duplex was buffer exchanged to a solution containing 150mM NaCl, 10mM NaMOPS and 50uM EDTA at pH7. The [DNA] for all experiments was 1 mM.

#### Mixer Design and Fabrication

The mixer portion of these devices is made of concentric capillaries; a central, sample-carrying fused silica capillary supply line (Polymicro Technologies, Phoenix, AZ) is held inside a larger glass tube (320 µm inner diameter, Polymicro Technologies, Phoenix, AZ) with custom Kapton centering spacers. The supply line is polished and beveled to a tip and an additional capillary, which acts as a mixing constriction and delay line, is held just downstream with a 50-100 µm gap. In this gap, the sample is flow-focused to a narrow stream and becomes fully sheathed by a $Mg^{2+}$ -containing buffer, which diffuses into the sample stream to initiate the reaction. With a single mixer, flowrates are varied to reach multiple timepoints, typically with a factor of two total range. The length and inner diameter of the mixing constriction is also changed to reach different timepoint ranges. A total of three different injectors were designed to reach timepoints from 6 ms to 1000 ms (Table S1).

The entire mixer is encapsulated in a larger glass shroud (750 µm inner diameter, Sutter Instrument), which is flame polished to create the nozzle opening. The end of the mixing constriction is held with another set of Kapton centering spacers, and as the liquid exits the capillary, it is exposed to a helium gas sheath (30 mg/min flowrate) which further thins and accelerates the stream to create a free-standing jet.

To achieve good signal-to-noise data for S/WAXS, it is crucial to have a pathlength that is as large as possible to maximize the scattering signal. Typical GDVN jets are only a few microns in diameter. To create thicker jets, closer to a target of a 10 µm diameter, the glass shrouds were flame polished to create wider openings of 110-150 µm, instead of the more typical 80-90 µm. Jet widths were measured 150 µm downstream of the nozzle opening at each flowrate condition with a liquid-jet imaging microscope equipped with backlight illumination and a Zyla CMOS camera (ANDOR, Oxford Instruments, Abingdon, UK; (*41*)). The overall nozzle geometry (tip of mixing constriction, shape of nozzle opening, distance between the constriction and the nozzle aperture) influence the size and stability of the jet, but generally wider apertures result in wider jets. Within a single nozzle, higher flowrates result in thicker jets. Additionally, since the sample is fully sheathed, only the center of the jet contains RNA. The following formula approximates the sample stream width:

$$d_{sam} = \frac{d_{jet}\sqrt{\dot{V}_{sam}}}{\sqrt{\dot{V}_{tot}}}$$

where $d_{sam}$ is the sample stream diameter, $d_{jet}$ is the total jet width for that flow condition, $\dot{V}_{sam}$ is the volumetric flowrate of the sample, and $\dot{V}_{tot}$ is the total volumetric flowrate (sample and buffer sheath combined). When designing the mixers, the sample flowrate was maximized to give as wide of a sample stream as possible, while balancing timing dispersion and sample consumption concerns. As the sample flowrate increases, it creates a wider sample stream, which increases the uncertainty in the measurement as the $Mg^{2+}$ ions need to diffuse further for complete mixing. These ions, however, diffuse extremely quickly, so timing dispersion concerns were mostly limited to the 6 ms and 10 ms timepoints. Table S2 details the jet width for each measurement.

**Data Collection and Real-Time Analysis at CXI**

*Acquiring the time series at CXI*

The initial state (0 ms, before mixing) of the RNA was taken with a Mg-free buffer sheath of 1 mM NaMOPS, 2 mM NaCl, 50 μM EDTA at pH 7. Similarly, the scattering profile of an AT DNA duplex was acquired with a Mg-free buffer sheath of 150mM NaCl, 10mM NaMOPS and 50uM EDTA at pH 7. Time-resolved data were acquired from RNA at either 1 mM or 0.5 mM in 1 mM NaMOPS, 2 mM NaCl, 50 μM EDTA at pH 7. For mixing experiments, the sheath contained 1 mM NaMOPS, 2 mM NaCl, 50 μM EDTA, 25 mM $MgCl_2$ at pH 7 for [RNA]= 1 mM experiments, or 1 mM NaMOPS, 2 mM NaCl, 50 μM EDTA, 15 mM $MgCl_2$ at pH 7 for [RNA]=0.5 mM experiments. Background profiles for subtraction were acquired by flowing Mg-free buffer in the sample stream and the appropriate $Mg^{2+}$ containing buffer in the sheath. High pressure reservoirs (1.2 mL for sample,Neptune Fluid Flow Systems, LLC., Knoxville, TN), 10-40 mL for buffer (KNAUER Variloop, KNAUER Wissenschaftliche Geräte GmbH, Berlin, Germany)) and home built switch boxes (using VICI valves, Valco Instruments Co. Inc., Houston, TX) facilitated an easy transition between sample and buffer.

Sample present and sample absent profiles were collected in succession. The RNA absent sample (buffer background) was acquired either immediately before or after the RNA present sample. In the best case, no adjustments were made at any time during these two experiments. Measurements at different [RNA] were interleaved to reduce nozzle changes. For the case of buffer background before sample data collection, a typical experiment proceeds as follows. The sample line was set to the Mg-free buffer, and the appropriate Mg-containing buffer was loaded into the sheath line. Flow rates were set for the desired time point. Once the jet stabilized, as assessed optically and with real time X-ray data analysis feedback with OM (see below), 5 minutes of data were collected. The sample stream was then switched to flow RNA, and flowed for about 5 minutes to purge the ~2 meter long supply line (necessitated by the CXI vacuum chamber design). The presence of RNA was assessed by the appearance of a visible scattering signal on the detector. Five minutes of data were collected from this point. After acquisition of the sample-present point, the flowrates were adjusted for the next timepoint, and another 5 minute acquisition began for the sample measurement of the next dataset. Following acquisition, the sample stream was switched to the Mg-free buffer and another 5 minute transition was allowed to clean the supply line. Subsequently, the buffer match was collected for 5 minutes. This buffer, sample, sample, buffer pattern allowed for maximum efficiency by minimizing buffer/sample transitions, while ensuring accurate buffer background matches. As discussed below, data quality depends on acquiring the buffer/sample sequence while flow conditions and beam position are as identical as possible.

Key to ensuring smooth data collection was the use of OnDA (online data analysis) Monitor (OM) (*46*). After subtraction of the detector dark image, each image was analyzed by an azimuthal integration around the beam center. Masks were created to remove spurious scattering flares from the jet as well as inactive pixels. Different masks were created for each run, as the jet widths were not constant, as discussed in Table S2.

This software provided real time scattering profiles, which helped judge jet stability and when supply lines were purged for buffer to sample transitions. Jet stability using the OM live-feedback software was assessed by visualizing radial stacks of WAXS profiles within seconds of X-ray illumination, monitoring continuity of the diffuse water peak at high-q values as well as stability of the overall scattering intensity.

*Background subtraction and acquiring solution scattering data at an XFEL*

Accurate solution scattering experiments require subtraction of a background (sample absent) scattering profile from a sample present scattering profile. Under normal conditions, where a fixed path sample cell is used (as is the case for all of the synchrotron and lab source data reported here), both profiles are acquired from samples that are sequentially loaded into the same sample holder. Corrections for any variations in beam intensity are made following standard practices, which usually involves scaling by the transmitted beam intensity (*47*). However, at the XFEL, the solutions are flowing in liquid jets whose size depends on the details of the injector design as well as the flow rates. In the absence of rarely occurring nozzle freezing events, the jet size remained stable for each sample condition, assessed via optical monitoring and using OM to view WAXS profiles and total scattering intensity. We developed the following protocol to perform accurate background subtraction from profiles acquired in flowing jets at the CXI beamline.

As a first step, it was important to assess the contribution to the scattering profile from the beamline alone (the beamline background). Because of the vacuum environment at CXI, the scattering from the chamber, and from the nozzle inside the chamber with its helium sheath, are more than a thousand times smaller than the scattering from the buffer jet (Figure S4), so this contribution is safely neglected.

To perform accurate buffer background requires careful normalization of each measured profile. Each profile must be individually scaled to account for all sources of variation. Three critical parameters were required to perform accurate background subtraction. We had to account for: the changing x-ray pulse intensity, variations in the sample thickness, to which the WAXS signal is directly proportional, as well as the sample temperature at the intersection point. We consider each of these factors in turn.

First, because of the stochastic way in which XFEL pulses are generated, pulse to pulse variations in energy are expected; most pulses fall within a factor of 4 of the mean intensity. Because the strength of the scattering is proportional to the pulse intensity that generates it, each profile must first be normalized by the intensity. For most solution scattering experiments, the profile is normalized to the transmitted beam intensity; however, because the flowing jet is so thin, transmission is nearly 100%. Thus, we recorded the intensity of each pulse using a Wave8 monitor which consists of a thin Si3N4 target and 8 photodiodes that collect backscattered X-rays. The X-ray pulse intensity is linearly proportional to the sum of the diode intensities. The beam position is calculated by the relative intensities of the 8 diodes (*48*). This value was recorded for each trace, and each profile was normalized to account for variations.

Second, a fixed path length is ideally maintained in solution scattering experiments. Typically, a fixed thickness sample holder contains the sample, however our liquid jets are freely flowing. Thus, we must account for the jet's thickness at the point where it intersects the beam, especially because the jet naturally thins out as it approaches its droplet breakup region and because there can be small motions of the beam, resulting in illumination of a different spot on the jet (either at a different height or a different width, which can result in being off center of the jet). After each profile is scaled to account for variations in XFEL pulse intensity, it was subsequently scaled to account for sample thickness. We chose to scale each profile to the integrated intensity of the first water peak. With the water peak intensity maximum located near q = 1.9 Å $^{-1}$, we integrated the profile over a q range from 1.6 to 2.25 Å$^{-1}$.

To validate this method for normalizing with sample thickness, we tested it at NSLS II because it assumes that the nucleic acid scattering is insignificant when integrated over this region. We acquired WAXS data on the single strands (rU30 and rA30) in a cell with a fixed sample thickness. Data were normalized either using the traditional approach (transmitted beam monitor) or by scaling to the integrated intensity of the water peak. Figure S5 shows that both normalization methods yielded results that are the same within noise fluctuation, suggesting that scattering derived from the RNA macromolecules is negligible in the q~2.0 Å $^{-1}$ regime.
Finally, proper background subtraction requires that the sample-present and sample-absent profiles are acquired at the same temperature. Because the liquid jet is flowing into the CXI vacuum chamber it is rapidly supercooled (*49, 50*) at a rate that depends both on jet speed (in this case ~10-50 m/s), thickness, and distance from nozzle. Evaporative cooling of the water occurs at rates of up to $10^7$ K/s under these conditions (*49, 51*). Temperature has a dramatic effect on water scattering, it alters the water structure factor throughout the curve. The water temperature can be readily ascertained by measuring the position of the first water peak near q = 2 Å $^{-1}$, which reports on the structural ordering of water (*52*). Measurements of I(q) at room temperature, acquired on our lab source, locate this peak at q=2.05 Å$^{-1}$ at 25˚C, and q= 2.03 Å$^{-1}$ at 9˚C. At LCLS, the peak is found near q=1.9 Å $^{-1}$. Comparing this to values from (*49*), which report the peak of the structure factor, we estimate that our sample temperatures are just below 260 K, in the supercooled regime. Temperature also affects the water scattering profile at lower q. Here, the detailed shape of the scattering profile reflects changing thermodynamic parameters of the water including isothermal compressibility and correlation length (*52*). Supporting information from Ref. (*49*), plotted and shown here as Figure S6, illustrates the effects of changing temperature on the scattering profiles throughout the q region of interest to this work. We plot the water structure factor from that work at temperatures of 258 and 251 K. Changes in this lower q region are particularly significant for our experiment, as they may be comparable in size to the time-resolved signals we seek to measure.

As part of our data analysis algorithm, we first selected profiles whose water peaks fell within a given q range for each run. We imposed an additional requirement to limit variations in temperature through the run, perhaps resulting from relative motions of the beam and the jet. We integrated the intensity in the low q portion of the curve, (from 0.3 to 0.9 Å$^{-1}$, a region where we detected variations in our control samples), and divided this sum by the integral over the water peak, discussed above. This ratio was computed for each beam and water peak height normalized profile, and over the course of a run, traced out a nearly gaussian curve. We selected curves for further analysis that fell within one sigma of the mean. Once this ‘temperature’ selection was made, the previously scaled azimuthal traces are summed and an average is computed to represents the either ‘sample present’ or ‘sample absent—buffer background’ trace for each condition. Note that if the nozzle was moved or replaced in between the buffer and sample collection, the data could not be properly scaled due to variation in temperature, and the point was discarded.

Once both the buffer and sample curves are thickness normalized and are acquired at a single temperature, the data point for each condition is derived by direct subtraction of the buffer from that of the sample under identical flow and nozzle conditions. Typical profiles are shown in Figure S7. Once the subtracted profile was obtained, other standard corrections were applied to account for solid angle in the WAXS regime, as well as beam polarization (*49*). These are the curves shown in Figure 3 of the paper. Offsets were applied for display purposes.

For one of the data points reported in the paper (t=60 ms), we acquired 36920/37503 pulses for the RNA present/RNA absent sample, 36192/36734 had acceptable pulse energies, 34372/35717 intercepted the jet, and 22511/27539 were similar in temperature. Overall, 71% of the data acquired for this point met our criteria, and this corresponded to between 3 and 4 minutes of data acquisition for this condition. Figure S8 illustrates this process as a flow chart.

#### Modeling connects features in the scattering profiles with molecular structures.

To aid in the interpretation of the peaks that appear at high q features, we turned to models. Beginning with a .pdb file that results from our WAXS-driven MD results of (*19*) for the long hairpin triplex in $Mg^{2+}$ we used WAXSIS (*53, 54*) to compute the scattering profiles out to q = 2 $Å^{-1}$. Results from these computations are shown in Figure S1. Panel a shows the profile computed from the full molecule, which is a 29 base pair hairpin interfaced with a 24 base triplex forming single strand, visualized using Chimera (*55*). The molecule is mostly in the triplex form, as shown to the right of panel a. The scattering profile was computed from a single frame of the simulation. Panel b shows the profile computed from the triplex piece of the full pdb file, extracted using Chimera. Note the appearance of four high q peaks near q=1.05, 1.4, 1.7 and 1.9 $Å^{-1}$. Following the protocol introduced in Ref. (*23*), the profiles of just the backbone molecules (phosphates plus sugar) were computed and are shown in panel c. Note that the first three of the above referenced peaks persist, while the fourth, highest q peak is absent. As noted in Ref. (*23*), peaks in this region reflect the arrangements of the phosphorous and oxygen atoms along the backbone. Panel d shows the scattering profiles of the bases alone, and the highest of the four peaks of panel b is identified with base stacking, as concluded in Ref. (*23*) for a DNA duplex. Finally, panel e shows the experimental profile of the hairpin triplex, from Ref. (*19*) deposited as SASDK26 in the SASBDB.

#### Low q changes.

Although this paper focuses on the high q correlations that report RNA structural motifs, data at lower q are also informative. They reflect the appearance of features with larger spatial dimensions in the sample. A comparison of the data from the three model structures of Figure 1 (the single strand, duplex, and triplex), shown here as Figure S2, highlights significant changes in the scattering profiles in the form of a broad shoulder that appears just above q=0.4$Å^{-1}$. Past machine learning studies of duplexes (*21*) associate this peak with the molecular radius, and it shifts between the duplex and triplex conformations. It is absent from the profiles of single stranded data, which do not have a radius defined by the interaction of multiple strands. Importantly this shoulder shifts to lower q as the molecule transitions from mostly duplex to mostly triplex, reflecting the increasing radius of the triplex. The prominence of the peak depends on the exact molecular conformations which are dynamic and depend on length salt and sequence. Changes in this q range are seen in the XFEL data and correlate with the appearance of higher q features.

#### Determining the proper [$Mg^{2+}$] level to initiate the reaction.

To find the proper amount of $MgCl_2$ required to initiate folding, we performed static titrations on our lab source, beginning with the low salt starting buffer and spiking in concentrated solutions of Mg in buffer. The goal was to determine the total Mg concentration required to fold the RNA at its different concentrations. At 0.5 mM RNA, the molecule appeared folded (assessed by examining the high q data, using a Kratky plot of $Iq^2$ vs. q), when the total Mg concentration equaled 5 mM. More Mg was required to fold the 1 mM RNA sample. In this case, nearly 10 mM of Mg (total) was introduced.

For mixing, we used a reaction threshold of 30%, meaning the sample was considered fully mixed when the concentration of $Mg^{2+}$ at the center sample stream reached 30% of the level in the outer, sheathing solution. Thus, for the lower [RNA], we considered the reaction initiated when the [Mg] reached 4.5 mM (30% of the 15 mM $MgCl_2$ sheath), and for the higher [RNA], we considered it initiated when [Mg]= 7.5 mM (30% of the 25 mM $MgCl_2$ sheath). It is plausible that the reaction was initiated at a slightly earlier time in the lower concentration sample, based on the increased ordering of the backbone visible at t=6ms. Data are shown in Figure S3.

#### The effect of temperature on scattering profiles of nucleic acids.

The high q scattering of nucleic acids can be strongly impacted by the nature of the solvent around the biomolecule. The scattering of bulk water is temperature dependent (Figure S6), and we must consider the temperature dependence of the hydration water around the biomolecule, whose interactions with the nucleic acid, or the ions around it (*56*), may be altered as a result of changes in the water structure. For example, it has been established that DNA has a chiral hydration spine (*57*), and the changing properties of liquid water, as described above, may alter the nature of this 'bound water', resulting in a temperature dependent change in the DNA scattering profile.

As a control, we measured the scattering profile of a model nucleic acid system, a DNA duplex consisting of a 25 nucleotide strand of A bases, that has been annealed with a 25 base strand composed of T bases. This A-T duplex displays numerous peaks in the high q region (*23*). To explore whether temperature affects the wide angle scattering profile of this A-T duplex, we performed experiments on the BioXolver lab source at fixed temperatures of 25˚C and 9˚C. Temperature dependent changes are detected both in the buffer scattering and in the DNA (as assessed after buffer subtraction). Figure S10 shows scattering of buffer acquired at 25˚C and at 9.4˚C. These profiles display a q dependent offset in the range $0.4 < q < 2.0$ Å$^{-1}$. Consistent with the above discussed reports of the temperature dependence of water scattering, made at both SSRL and LCLS (and shown in Figure S6), we measure changes in both the location of the water peak and in the scattering at lower values of q. Background subtracted profiles from the AT duplex, measured at both 9˚C and 25˚C are shown in Figure S11. Changes are detected above q ~ 0.3 Å. A WAXS profile acquired at LCLS, where the solvent is supercooled (water peak) is shown as the third line on the figure. A significant decrease is seen in the same q region, signaling a potential temperature dependent effect. This latter curve was acquired in a liquid jet from a mixing injector at CXI, though no mixing was triggered (there was no difference in the buffer around the DNA and the sheathing buffer).

Interestingly, the folded RNA trace (Figure S9) shows deviations when acquired at room temperature and at LCLS. In both cases, the profiles 'match' at low q and at the highest q, but deviate in a region that tracks with the length scales of duplex and triplex features.

#### RNA folding at reduced concentration: 0.5 mM.

Figure S12 shows the time series acquired with lower RNA concentration. Following the guidance of static $Mg^{2+}$ titration experiments (Figure S3), and with the mixing threshold set at 30% (Methods), the [$MgCl_2$] in the sheathing buffer is 15 mM when [RNA]=0.5 mM and 25 mM when [RNA]=1 mM. Because the exact composition of the ion atmosphere around the RNA is unknown, e.g. the number of bound vs. free Mg ions cannot be determined from this experiment (*58*), it is plausible that the free [Mg] is higher for this time series, resulting in slight variations in folding kinetics between the two conditions. Nevertheless, the order of assembly is the same. Duplex features appear by the first measured time point. At later time points, the major groove peak disappears, and the triplex backbone structure strengthens. At longer times, the base stacking peak appears, locking in the structure. These trends are exactly recapitulated in the lower q peak (0.4 $Å^{-1}$): the loose shoulder becomes better defined on time scales longer than 100 ms.

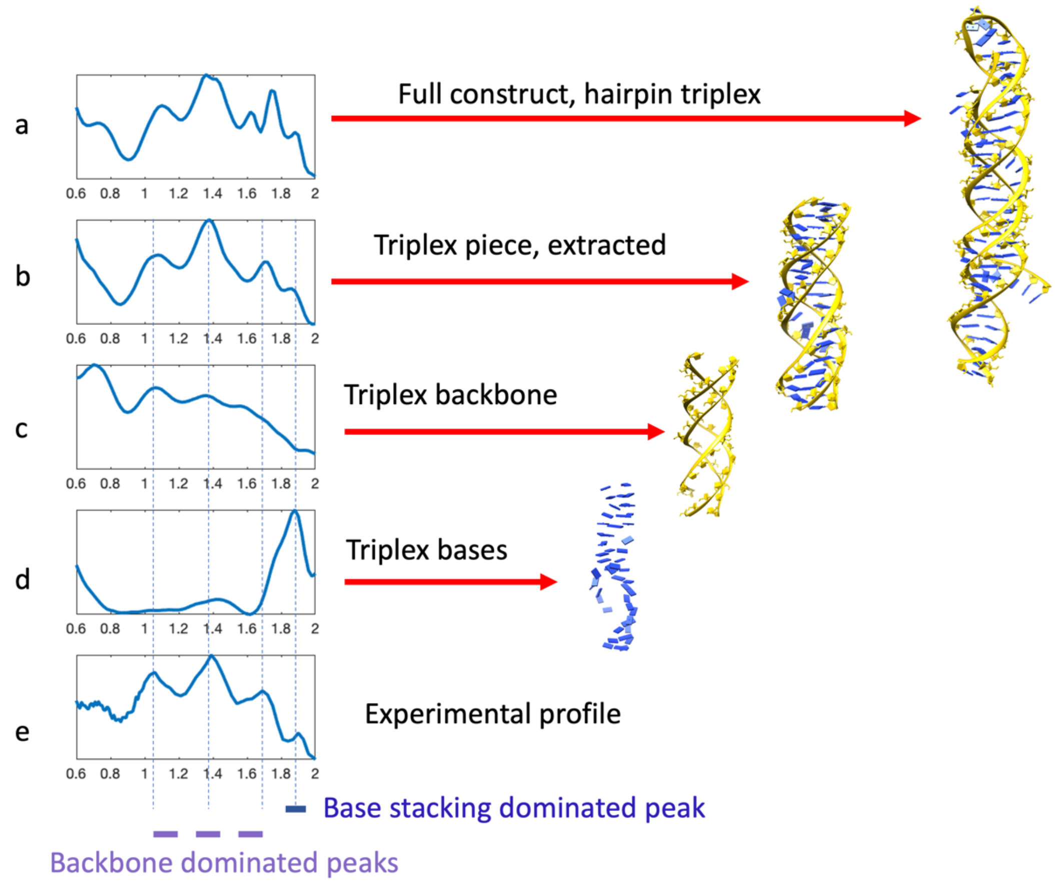


Figure S1 connects the features of triplex scattering profile with real space molecular structures. As described and referenced in the text, profiles are computed via WAXSIS. The model structure shown here is the one pictured in Figure 1 panel C of the main text.

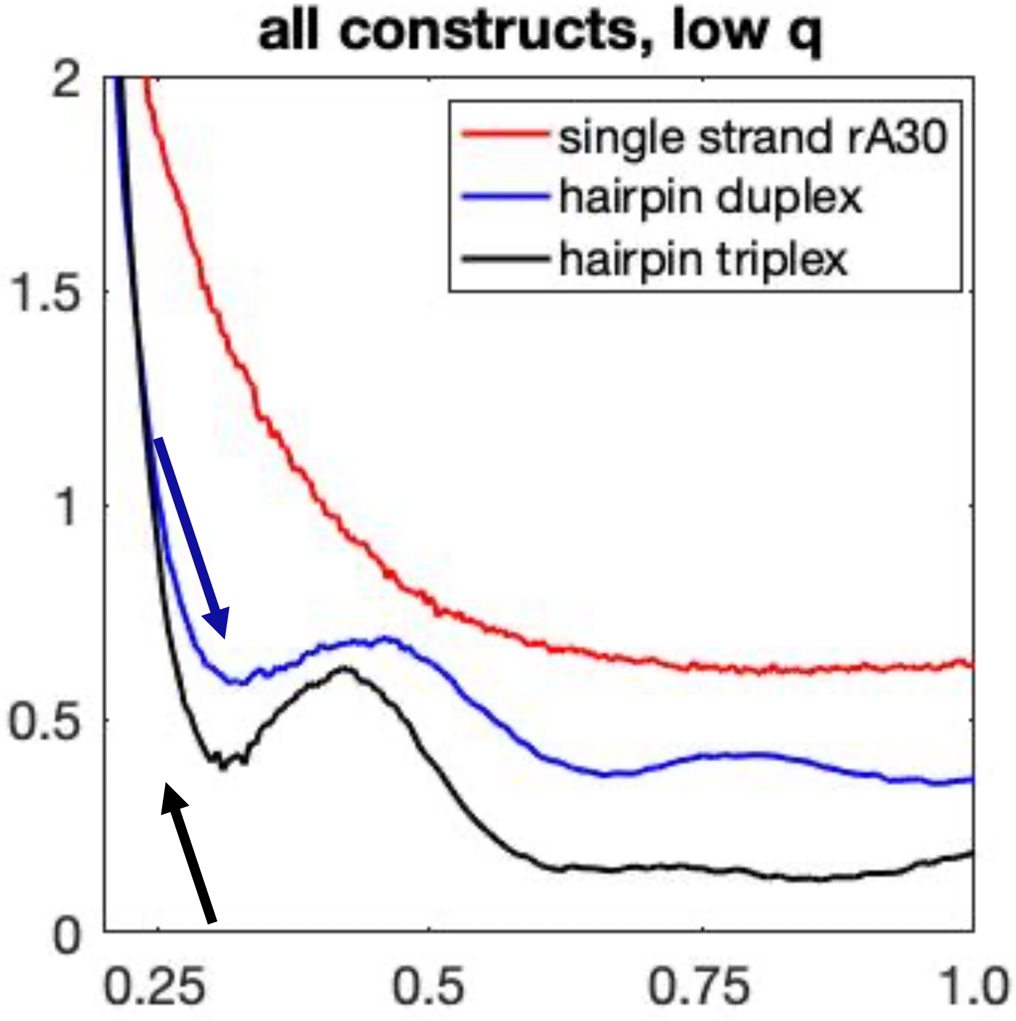


Figure S2 highlights low q features that reflect the molecular radius. Arrows indicate regions which change with molecular conformation. Scattering from an unstructured single strand is shown in red, from a helical duplex in blue and a triplex in black. These profiles were shown in Figure 1 of the paper, we simply emphasize a different q region here.

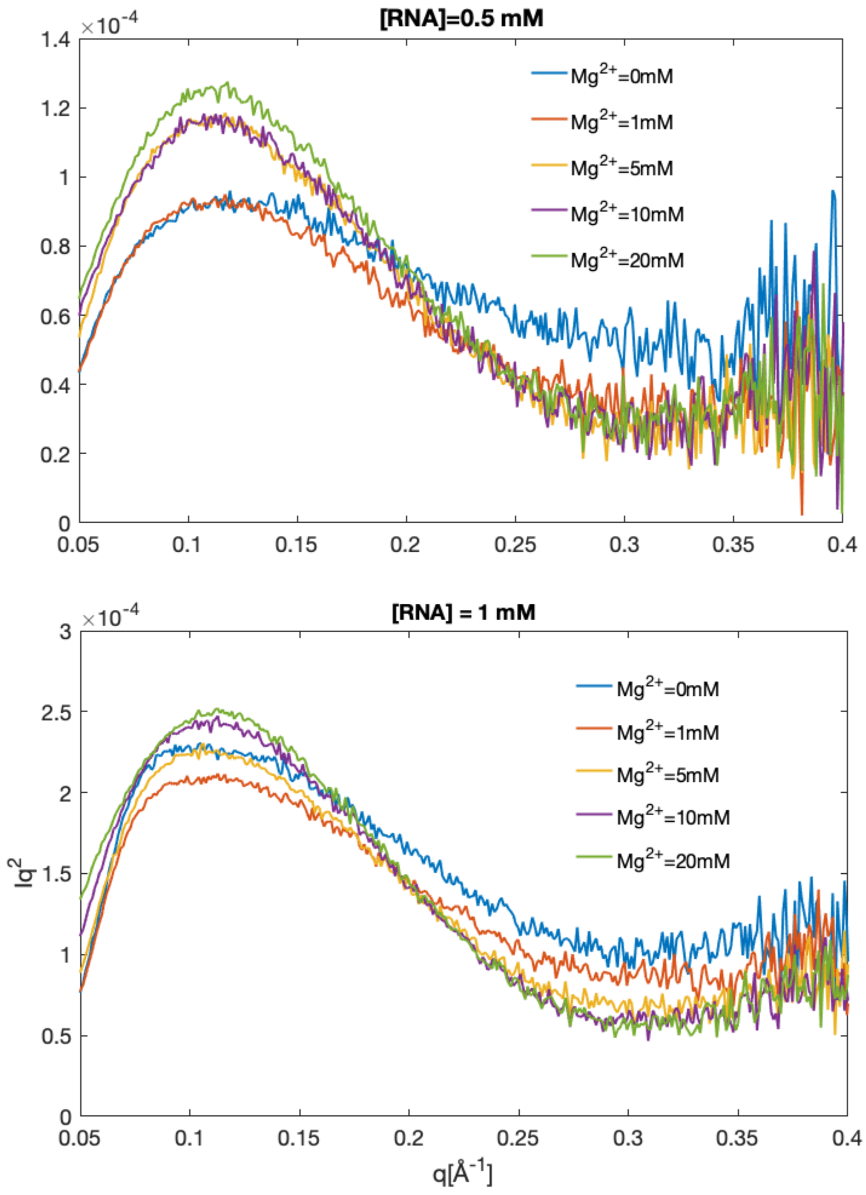


Figure S3 illustrates the $Mg^{2+}$ dependent folding of the UAU12 construct, showing static data with increasing [Mg] acquired on the BioXolver source. These profiles are displayed as Kratky plots of $Iq^2$ vs. q, which emphasize the changes that indicate folding. More folded molecules have better defined peaks near q=0.12 $Å^{-1}$ as well as lower values at large q. From these curves, we selected the concentrations of 4.5 mM and 7.5 mM as reaction initiation thresholds for [RNA] at 0.5 mM and 1 mM.

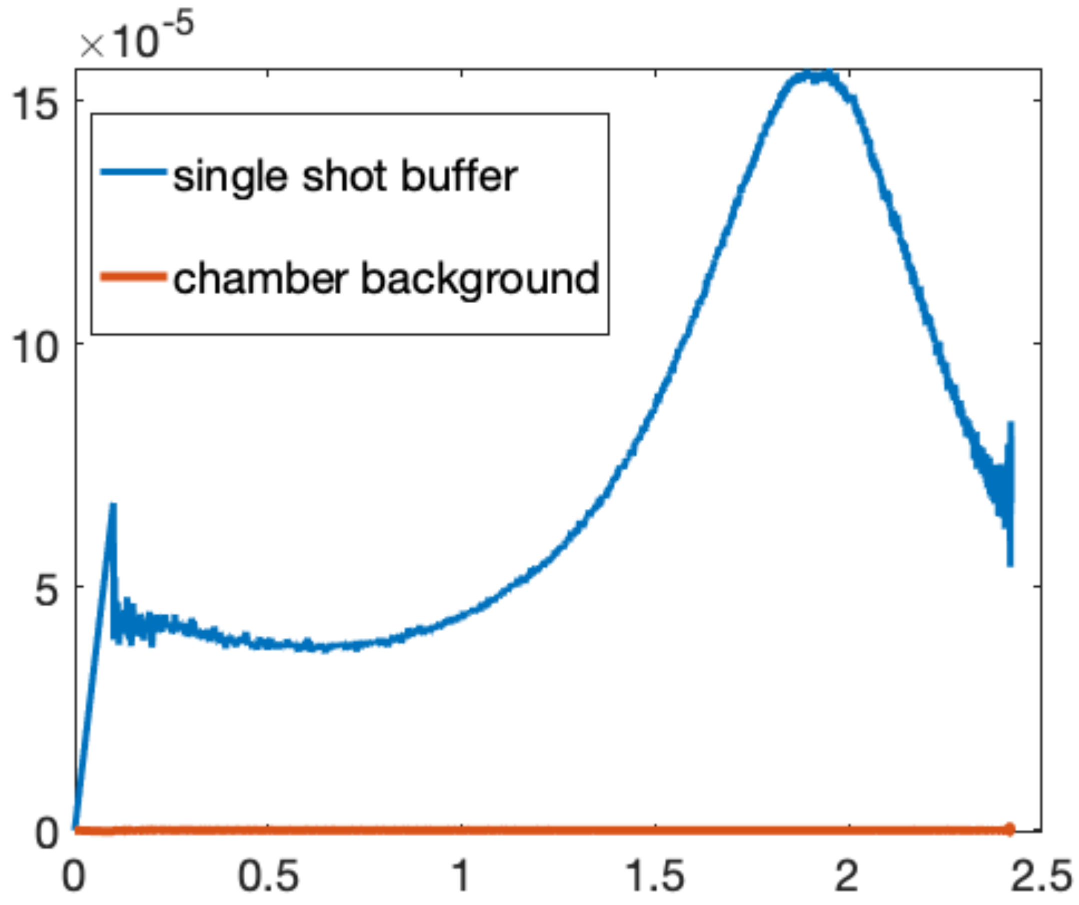


Figure S4 demonstrates the level of scattering the chamber background (brown), beam normalized, compared to a single shot of buffer scattering (blue) also beam normalized. The small contribution from the background justifies its omission from our analysis.

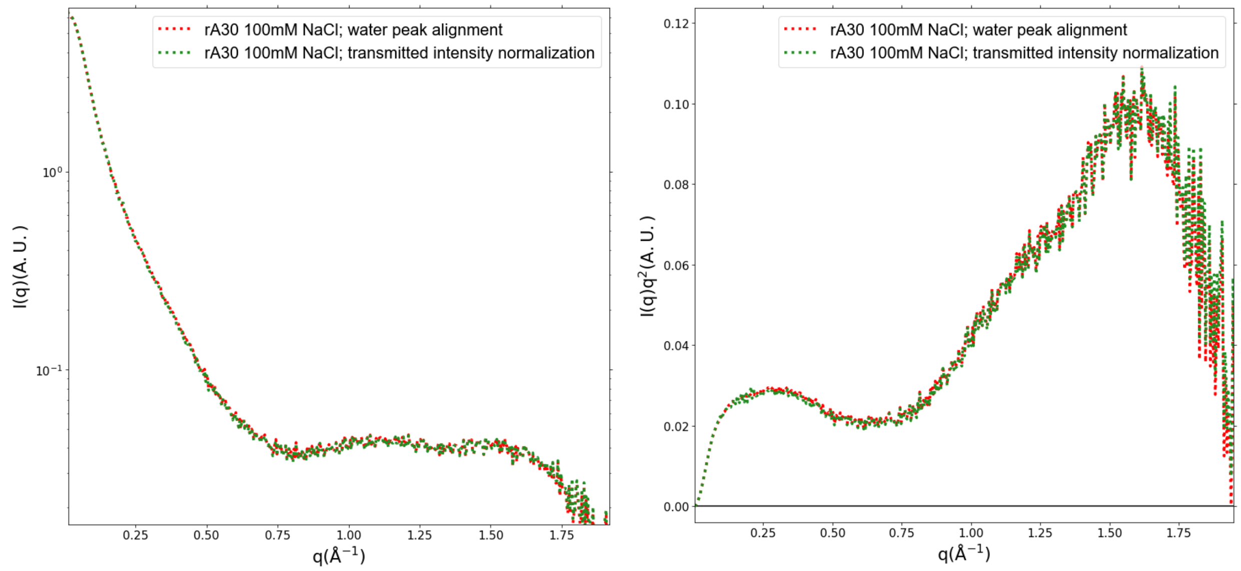


Figure S5 – Comparison of scattering profiles of rA30 in 100mM NaCl, where alignment of the sample and buffer scattering has been performed using alignment of the average water peak scattering intensity versus normalization by the transmitted X-ray beam intensity. Scattering intensities are plotted in arbitrary units. Data are plotted on semilog (left) and Kratky-transformed (right) axes to visualize agreement of the two methods in both low and high q regimes. All other single stranded RNA data showed similar levels of agreement.

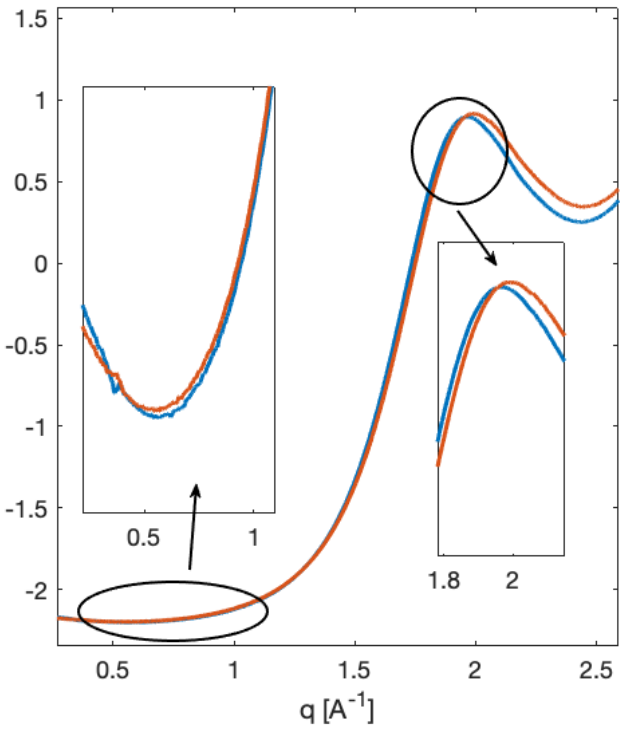


Figure S6 shows changes in the water structure factor, graphed from the supporting data of Ref. (*49*). These curves illustrate the temperature dependent changes that impact the shape of the curve. These curves were taken at T=258 K (red) and T=251 K (blue).

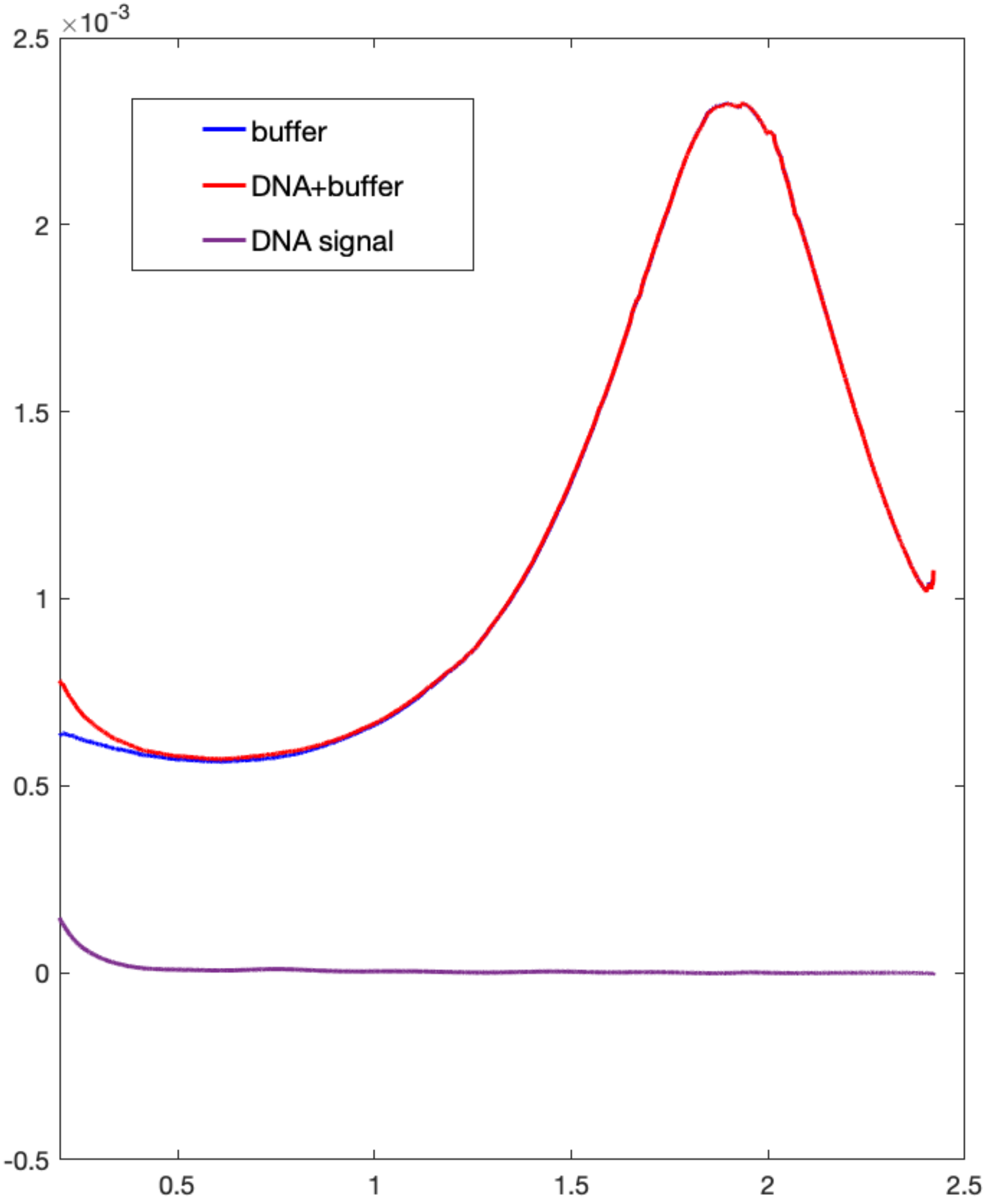


Figure S7 shows the beam and sample thickness normalized curves before and after subtraction. This plot shows beam and jet normalized buffer (blue), sample (red) and the difference (purple) from our control sample: the 25 base pair AT duplex. The subtracted curve is also shown in Figure S11.

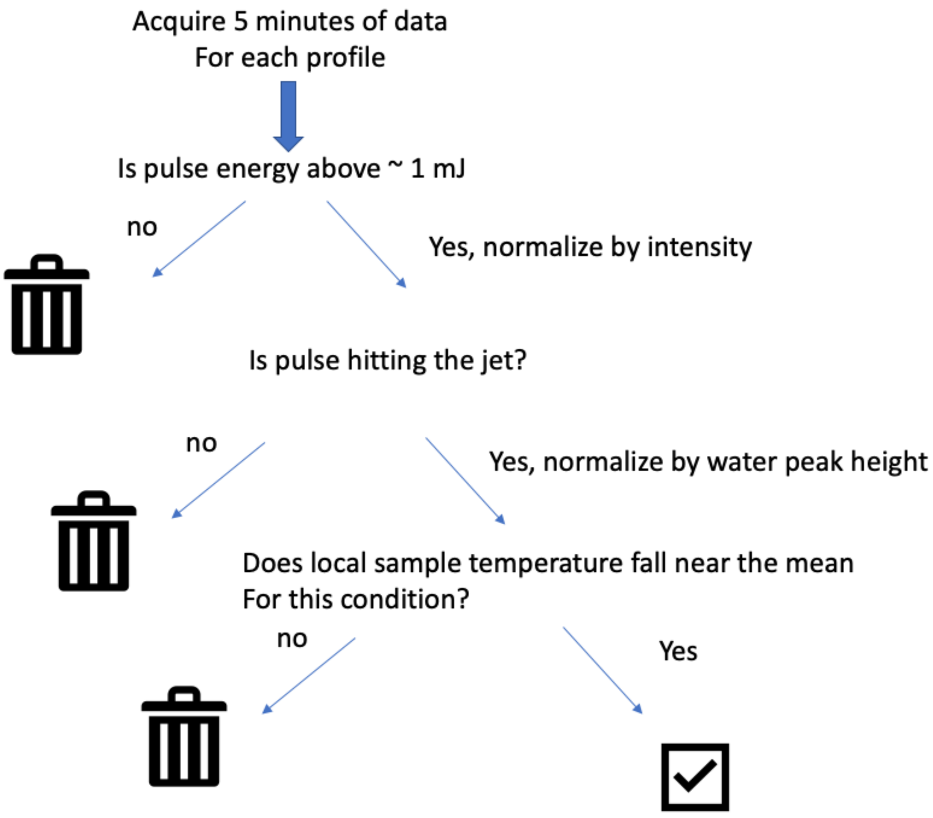


Figure S8 shows the pipeline for data analysis described in the text, as a flowchart.

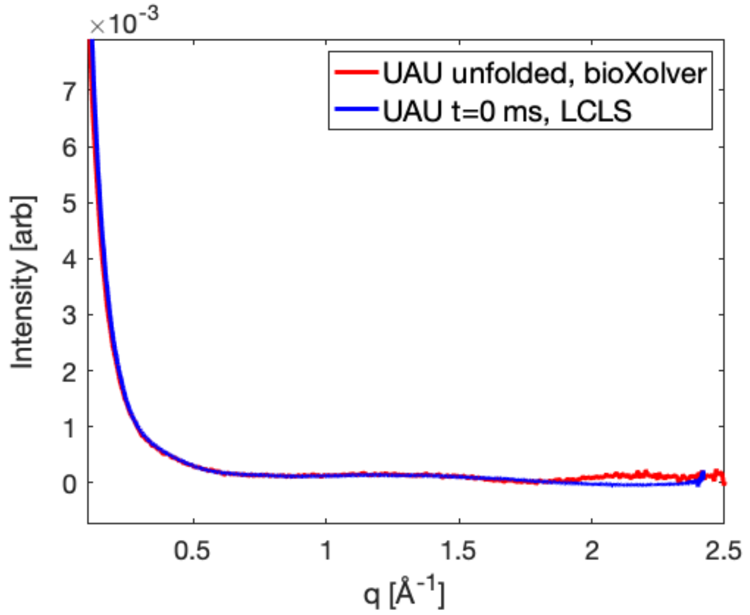


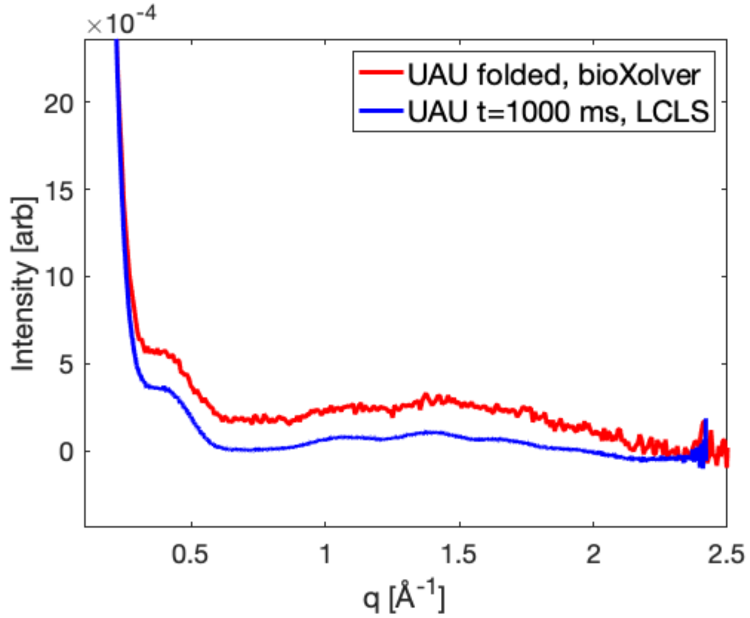


Figure S9 shows a comparison of both single stranded UAU12 and triplex UAU12, acquired at room temperature on our lab source using a thick sample (red, exposure time 120 seconds), compared to LCLS (blue).

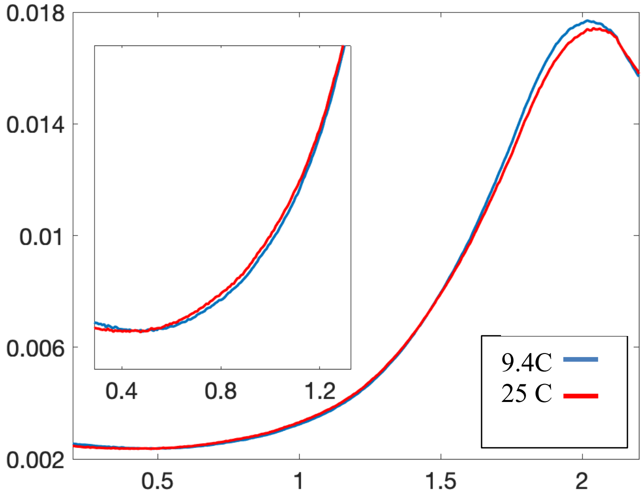


Figure S10 shows the temperature dependent buffer scattering, acquired on the BioXolver lab source, at two temperatures: 9 and 25 C. These trends, especially the q regions where there are changes, exactly echo the conclusions of Ref. (*49*).

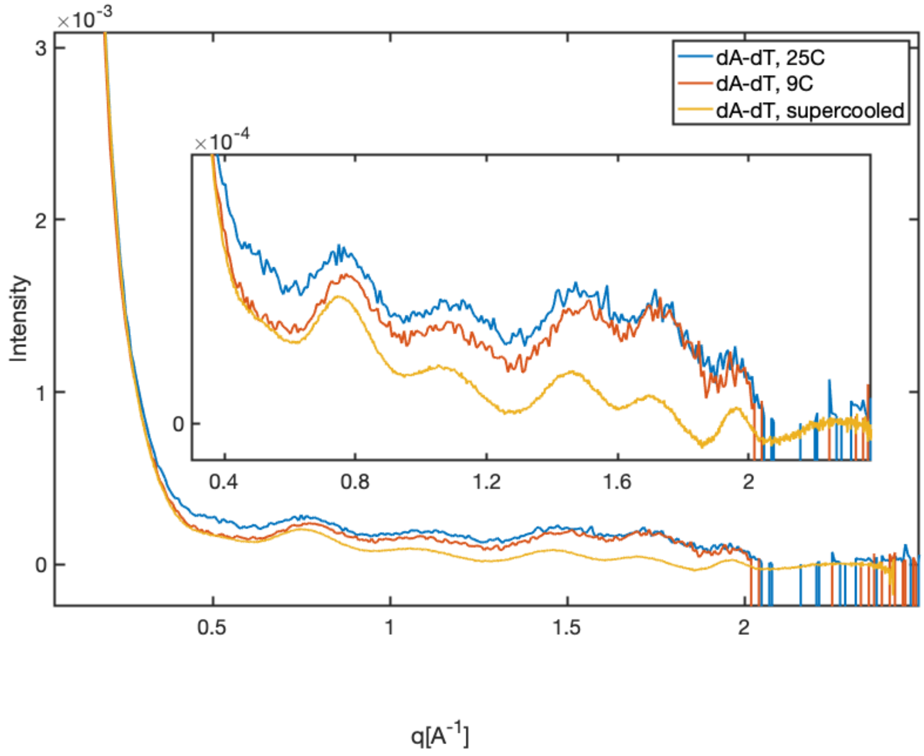


Figure S11 shows the temperature dependent scattering of our control molecule, a 25 base pair DNA duplex made from joining dT25 with dA25 (two 25 nucleotide strands). Note the deviations from room temperature to 9 C, and again, to the LCLS measured state at a lower temperature.

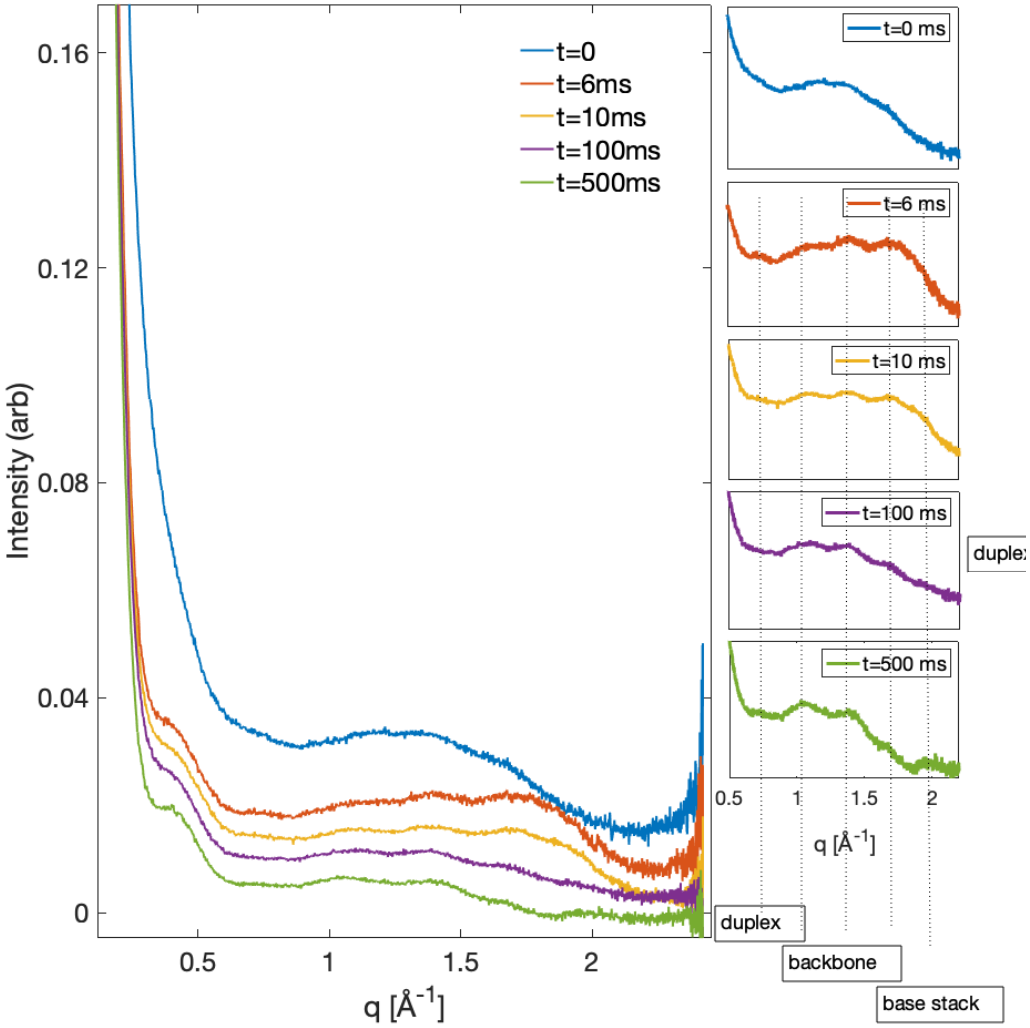


Figure S12 shows the time course of RNA folding for [RNA]= 0.5 mM. Here the sheath contains 15 mM Mg, and folding considered initiated when the [Mg] reaches 4.5 mM. The order of appearance of the peaks recapitulates the data shown in the paper for [RNA] = 1 mM. A small, major groove associated peak appears at the fastest measurement time, t= 6 ms. Other peaks, representing the triplex backbone also appear and stabilize at longer reaction times. A high q peak, reflecting base stacking becomes prominent by 500 ms, though it is visible at earlier times. Note also that the low q behavior reproduces what is measured at the higher [RNA]. The high q background varies between this series and the previous. This may be a result either of differences in [Mg], variations in temperature due to changing salt conditions, or smaller signal size, due to the reduction in concentration. and are the result of either temperature variation or the more challenged signal strength for the lower [RNA] data set.

Supplementary Tables:

| Timepoint (± uncertainty; ms) | Constriction Length (mm) | Constriction Inner Diameter (µm) | Sample Flowrate (µL/min) | Sheath Flowrate (µL/min) |
|---|---|---|---|---|
| 6 ± 1.5 | 17.8 | 50 | 9.6 | 123.3 |
| 10 ± 2.9 | 17.8 | 50 | 10.3 | 67.3 |
| 60 ± 19.1 | 32 | 100 | 19.1 | 72.3 |
| 100 ± 28.3 | 32 | 100 | 16.5 | 41.7 |
| 500 ± 34.3 | 25** | 100 | 20.6 | 80.4 |
| 1000 ± 96.6 | 25** | 100 | 18.8 | 34.2 |
| ** For timepoints 500 ms and longer, an additional delay stage is added after the constriction | | | | |

**Table S1:** Injector design and flow conditions for each timepoint

| Timepoint (± uncertainty; ms) | Nozzle Opening (µm) | Total Jet Width (µm) | Sample Width (µm) | Total Flowrate (µL/min) |
|---|---|---|---|---|
| 0 | 127 | 10.2 | 4.6 | 101 |
| 6 ± 1.54 | 90 | 6.4 | 1.7 | 132.9 |
| 10 ± 2.90 | 90 | 3.8 | 1.4 | 77.6 |
| 60 ± 19.12 | 90 | 7.1 | 3.2 | 91.4 |
| 100 ± 28.27 | 90 | 5.1 | 2.7 | 58.2 |
| 500 ± 34.3 | 127 | 10.2 | 4.6 | 101 |
| 1000 ± 96.6 | 127 | 6.4 | 3.8 | 53 |

**Table S2:** Nozzle openings and jet widths for each timepoint
